# Mob4 is essential for spermatogenesis in *Drosophila melanogaster*

**DOI:** 10.1101/2023.02.05.527206

**Authors:** Inês B. Santos, Alan Wainman, Juan Garrido-Maraver, Vanessa Pires, Maria Giovanna Riparbelli, Levente Kovács, Giuliano Callaini, David M. Glover, Álvaro A. Tavares

## Abstract

Gamete formation is essential for sexual reproduction in metazoans. In males, spermatogenesis gives rise to interconnected spermatids that have to differentiate and individualize into mature sperm. In *Drosophila melanogaster*, individualization of spermatids requires the formation of individualization complexes that synchronously move along the sperm bundles. Here, we show that Mob4, a member of the Mps-one binder family, is essential for male fertility but has no detectable role on female fertility. We describe the function of Mob4 during spermatid individualization, showing that Mob4 is required for proper axonemal structure and that the loss of Mob4 leads to male sterility associated with defective spermatid individualization and absence of mature sperm in the seminal vesicles. Transmission electron micrographs of Mob4^RNAi^ developing spermatids revealed defects in axoneme structure and abnormal mitochondria biogenesis. Importantly, we find that male fertility is impaired upon depletion of other STRIPAK components, suggesting that Mob4 acts through STRIPAK to support spermiogenesis. As we show that expression of the human Mob4 gene effectively rescues all phenotypes of Mob4 downregulation, the gene is not only evolutionary but also functionally conserved. We propose that Mob4 plays a role in regulating the microtubule- and actin-cytoskeleton during spermatogenesis. This study advances our understanding of male infertility by uncovering Mob4 as a novel gene required for sperm individualization.

## Introduction

Human infertility is a major global public health issue estimated to affect one out of six couples, with half of the cases arising from a male factor. The many cause of male infertility include defects in spermatogenesis or spermiogenesis. However, medical approaches to treat the underlying causes of reduced sperm production remain elusive. This is largely because of the limited knowledge of the molecular and cellular processes that regulate sperm production and its failure (Tuttelmann et al., 2018). As spermatogenesis is a highly conserved process, not differing appreciably between insects and mammals (Ramm et al 2014), *Drosophila melanogaster* offers a powerful model organism in which to uncover conserved genes that regulate spermatogenesis.

Adult male *Drosophila* have a pair of testes, with an apical end containing approximately 8–12 germinal stem cells (GSCs) (Hardy RW et al., 1979), each flanked by two somatic cyst stem cells (CySCs) (Gonczy P, DiNardo S, 1996) that eventually differentiate into a head cyst cell and a tail cyst cell, analogous to mammalian Sertoli cells. GSCs divide asymmetrically in the cysts to give rise to spermatogonial cells. Spermatogonia then go through four rounds of mitosis, generating 16 primary spermatocytes. Following an extended G2 phase of growth, the primary spermatocytes undergo two meiotic divisions to yield a cyst of 64 syncytial haploid spermatids, each (reviewed in Fuller MT, 1998). Spermatids being inter-connected by cytoplasmic bridges (Lin H et al., 1994; Hime GR et al., 1996). During post-meiotic spermatid differentiation, syncytial cysts of 64 haploid spermatids undergo synchronous differentiation, which requires formation of flagellar axonemes and acrosomes, remodelling of mitochondria and nuclei, and the polarization of elongating cysts and the plasma membrane (Tates AD, 1971; Tokuyasu KT, 1975; Lindsley DL, Tokuyasu KT, 1980). The fully elongated syncytium of 64 spermatids then undergoes a membrane remodelling process known as individualization as it generates individual sperm. Individualization begins with the formation of actin investment cones around each of the 64 needle-shaped nuclei (Fuller MT, 1998). The 64 actin cones within a cyst assemble into a structure referred to as the individualization complex (IC) that moves up the testes while the cyst membrane is remodelled and intercellular bridges are resolved to encase each sperm cell in its own plasma membrane (Fabrizio J et al., 1998). The IC moves processively along the spermatid bundle, stripping away unnecessary organelles, mitochondrial DNA, and cytoplasmic components, forming the dilation of the cyst known as the “cystic bulge” (Tokuyasu KT et al., 1972; Noguchi T, Miller KG, 2003). The cystic bulge is then detached at the end of the spermatid bundle where it becomes the “waste bag”. The organization of the actin cones and their movement as a cohort is critical for spermatid individualization. Cytoskeletal regulators, such as myosin V, myosin VI, cortactin, and Arp2/3 complex, have been shown to influence formation of actin cones and the synchronous movement of the IC (Hicks J et al., 1999). In myosin V mutants, for example, fewer actin cones form (Mermall V et al., 2005). Myosin VI acts to stabilize the actin cones and the Arp2/3 complex is required for the formation of the actin meshwork (Noguchi T et al., 2006; Noguchi T et al., 2008). Cortactin co-localizes with Arp2/3 complex and myosin VI at the leading edge of the actin cone during IC movement.

The individualization process is as a caspase-dependent apoptosis-like event, requiring functional proteasomes and membrane trafficking. Multiple *Drosophila* caspases and caspase regulators participate in this process and mutation of any of these genes leads to individualization defects.

The Striatin-interacting phosphatase and kinase (STRIPAK) complex is an evolutionarily conserved supramolecular complex with diverse functions in cell proliferation, migration, vesicular transport, cardiac development, and cancer progression (Hwang J, Pallas DC, 2014; Madsen et al., 2015; Neisch et al., 2017; Shi et al., 2016). In addition, several STRIPAK components participate in dendritic development, axonal transport, and synapse assembly (Chen et al., 2012; Li et al., 2018; Schulte et al., 2010). In *Drosophila*, Strip (a homolog of mammalian Strip1/2) is essential for axon elongation by regulating early endosome trafficking and microtubule stabilization.

STRIPAK complexes contains multiple components, some of which are mutually exclusive. STRIPAK integrates a variety of regulatory proteins that can direct the PP2A catalytical and regulatory A subunit to specific targets (Shi Z et al., 2016; Ribeiro P et al., 2010; Virshup DM, 2000). In *Drosophila* and mammals, a STRIPAK-PP2A complex containing Mob4 and Cka was reported to inhibit Hippo signalling (Zheng Y et al., 2017; Couzens A et al., 2013; Ribeiro P et al., 2010). STRIPAK complexes acts as platforms to integrate upstream inputs to the Hippo pathway (Bae S, 2017; Chen R, 2019). One such example is the regulation of MST2 (a member of the mammalian Ste20-like kinases, mammalian homolog of Hippo) by the STRIPAK component SLMAP (Sarcolemmal Membrane-Associate Protein). SLMAP has been shown to recognize phosphorylated MST2, thereby recruiting MST2 to STRIPAK for dephosphorylation and inactivation in rapidly dividing cells (Bae S, 2017; Zheng Y, 2017). It has been shown that members of the germinal centre kinase III (GCK III) subfamily of the Ste20 superfamily of kinases participate in STRIPAK complex and function to suppress the Hippo pathway in *Drosophila*, and more recently in humans (Bae S, 2020).

Mob4 can inactivate the Hippo pathway as a core component of STRIPAK complex, resulting in enhanced expression of growth-promoting genes. Moreover, a recent report revealed that in mammals, Mob4 forms a complex with MST4 kinase and disrupts MST1-Mob1 complex and increases YAP1 activity, which implies that Mob4 can be oncogenic (Chen M et al., 2018). In *Drosophila*, Mob4 is described to regulate normal synapse formation (Baillat G et al., 2001) and to be required for both centrosome separation and focusing of K fibers (Trammell A et al., 2008). Within the STRIPAK, Mob4 acts as molecular glue to tether the WD40 domain of STRN1 (mammalian homolog of Cka) to the main body of STRIPAK. In fact, point mutations targeting 3 residues that serve as phosphor-threonine binding sites to Strip1 (mammalian homolog of *Drosophila* Strip), decrease the formation of STRIPAK which led to deregulated Hippo signalling in cultured cells (Jeong BC et al., 2021). Nevertheless, although Mob4 is an integral part of the STRIPAK core, the molecular function of Mob4 remains poorly defined.

Here, we report an essential role for Mob4 in spermatogenesis in *Drosophila melanogaster* and show that the human Mob4 homologue can rescue the *Drosophila* phenotypes, indicating conserved function.

## Results

### Lack of Mob4 causes lethality and male sterility in Drosophila melanogaster

To study the potential functions of the Mob4 gene, we generated two different Mob4 mutant alleles. The first, Mob4^P^, was generated by imprecise excision of the P-element from the line KG4509 (see *Methods*). Mob4^P^ lacks most of the 1st exon, including the ATG start codon (Figure 1A). When homozygous, Mob4^P^ animals died during the third larval instar stage, although a small fraction (0.4%) of “escaper” flies reach adulthood. Such female escaper flies are fertile but escaper males are sterile. Such sterile males were able to mate normally but were unable to fertilize eggs. The second Mob4 mutant allele was a null that we generated using the CRISPR/cas9 genome editing system (Port et al., 2014). This null allele, Mob4^SVC^, lacks the entire Mob4 coding region (Figure 1A) and also showed 3rd instar lethality when homozygous or heterozygous to a deficiency uncovering Mob4. We were able to rescue the lethality of Mob4^SVC^ flies by ubiquitously expressing a wild-type GFP-Mob4 transgene, indicating that the lethality is a consequence of the mutation in Mob4.

**Figure 1.**
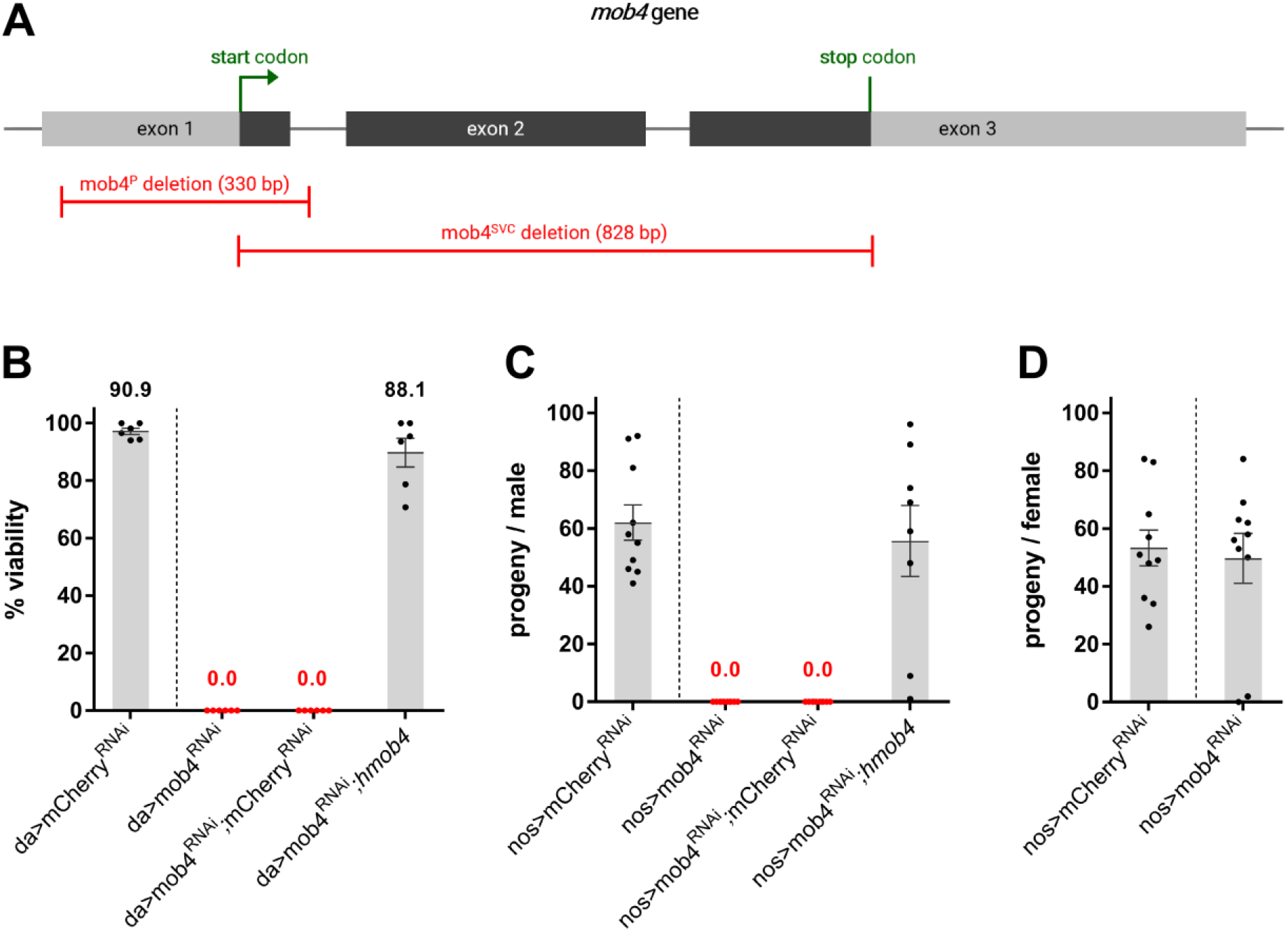
Mob4 is essential for viability and male fertility in *Drosophila melanogaster*. **(A)** Schematic of mob4 gene organization and CRISPR/Cas9-targeted mutagenesis to show the resulting deletion allele (red), mob4^SVC^. The mob4^P^ deletion allele was generated by imprecise excision of the P-element in the line KG4509. The endpoints of deletion in mob4^SVC^ and mob4^P^ were determined by sequencing and are indicated in basepairs (bp). **(B)** Viability of control, mob4^RNAi^ and rescued flies (mob4^RNAi^ and human mob4 (*hmob4*), both expressed under UAS-control in da-GAL4 flies). Percentage of viable flies was determined from number of eclosed adults relative to the number of pupae. **(C)** Male fertility test of control, mob4^RNAi^ and rescued flies (mob4^RNAi^ and *hmob4*, both expressed under UAS-control in nos-GAL4 flies). Males of each genotype were individually mated with wild-type females. Data points represent numbers of progeny from individual males. **(D)** Fertility test of control and mob4^RNAi^ females. Females of each genotype were individually mated with wild-type males. Data points represent numbers of progeny from individual females.

Intrigued by the sterility of male escapers, we sought to analyse the role of Mob4 during spermatogenesis. To this end, we chose to downregulate Mob4 in the male germline in two different UAS-Mob4^RNAi^ transgenic lines whose expression was driven by GAL4 driver lines with specific spatio-temporal expression patterns in fly testes (White-Cooper H, 2012). We used two different driver lines: nos-GAL4, which induces target expression in GSCs and spermatogonia, and bam-GAL4, which does not induce expression in the GSCs but in late spermatogonia and early spermatocytes. In this way, we could overcome Mob4 lethality and study Mob4’s role in male fertility, obtaining identical results for both UAS-Mob4^RNAi^ lines. When we used either the nos-GAL4 (Figure 1C) or the bam-GAL4 (data not shown) to drive Mob4 knockdown and crossed the resulting progeny with wild-type virgin females, none of the eggs hatched, recapitulating the mutant escaper phenotype. When we downregulated Mob4 in the female germline, there were no effects on female fertility (Figure 1D). When we used the daughterless (da)-GAL4 driver and the UAS-Mob4^RNAi^ reporter lines to achieve ubiquitous depletion of Mob4, this phenocopied the Mob4^SVC^ null mutant phenotype, resulting in complete lethality at the late larval/pupal stage (Figure 1B). Together, these results indicate that Mob4 is required for male fertility, but not for female fertility.

To ensure that the consequences of Mob4 RNAi downregulation were not off-target responses, we determined whether the Mob4^RNAi^ phenotype could be rescued by coexpression of the human Mob4 gene (UAS-*hMob4*), which is resistant to dsRNA directed against the *Drosophila* gene. In fact, we found that all phenotypes of Mob4 depletion were rescued by provision of human Mob4. Specifically, viability (figure 1B) and male fertility (figure 1C) were restored to control levels, indicating that the lethality and male sterility phenotypes upon Mob4^RNAi^ are specific to the depletion of the conserved gene Mob4.

### Mob4 is required for spermatid individualization

To explore the mechanisms underlying the sterility of males lacking Mob4, we compared testes from control young un-mated males with Mob4^RNAi^ males. There were no major morphological defects in Mob4^RNAi^ testes leading us to visualize spermatid axonemes by immunostaining with anti-pan polyglycylated tubulin antibody (Axo-49). In control testes, axonemes can be observed throughout the testes length (elongating cysts), in the terminal epithelium (sperm coiling) and inside the seminal vesicles (where mature sperm is stored until mating) (Figure 2A). By contrast, cysts of Mob4^RNAi^ testes were fully elongated but the seminal vesicle remained empty (Figure 2E). In addition, most of the Mob4^RNAi^ sperm bundles accumulated in the terminal epithelium (highlighted in pink, Figure 2E) and showed increased tail coiling, suggesting defects in sperm individualization.

**Figure 2.**
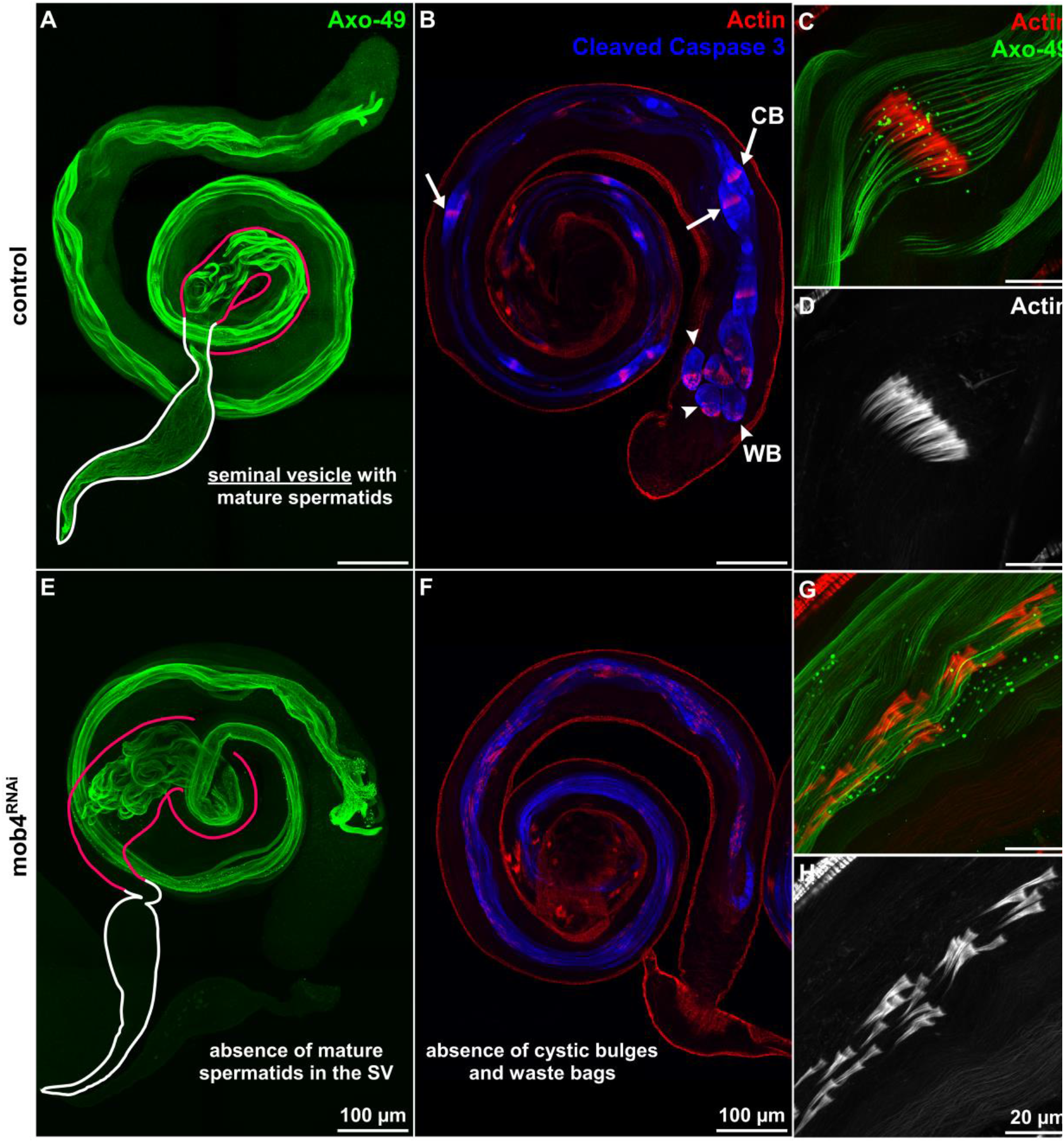
Individualization defects in Mob4 depleted testes. Control **(A)** and Mob4 depleted **(E)** testes from 2-day old males were immunostained with anti-polyglycylated tubulin (green) to visualize spermatid axonemes. The seminal vesicle (SV) and terminal epithelium (TE) are highlighted in white and pink, respectively. Control **(B)** and Mob4 depleted **(F)** testes were stained with phalloidin-594 and immunostained to reveal cleaved caspase 3 (blue). Cystic bulges (CB, white arrows) and waste bags (WB, white arrowheads) resulting from spermatid individualization can be seen in control testes **(B-D)** but are absent from mob4-depleted testis **(F-H)**. Actin cones and spermatid bundles from control **(C-D)** and mob4 depleted testes **(G-H)** visualized with phalloidin-594 and immunostained to reveal polyglycylated tubulin (green).

Spermatid individualization occurs in the final stage of spermiogenesis (Lindsley DL et al., 1980; Fuller M, 1993) in elongated spermatid bundles. It is mediated by individualization complexes (ICs), which contain a cluster of 64 actin-rich structures, known as actin cones. During sperm individualization, the IC translocate along the length of the cyst and remove excess cytoplasm and organelles into so-called ‘cystic bulges’ (Figure 2C-D). Concomitantly, spermatids become sheathed in plasma membranes and become individualized. By the end of the individualization process, the 64 actin cones are degraded together with the syncytial cytoplasm in a structure known as the waste bag (WB) (Noguchi T et al., 2006; Fabrizio J et al., 1998). Disturbances of any of these processes may lead to individualization defects and the stagnation of sperm bundles in the terminal epithelium.

To investigate potential requirements for Mob4 during the individualization process, we used immunofluorescence to examine the formation of actin cones in control and Mob4^RNAi^ testes. In control testes, actin cones move uniformly along the sperm bundles forming the cystic bulge (Figure 2C-D). However, in Mob4^RNAi^ testes, the ICs were scattered along the sperm bundles (Figure 2G-H).

Defects in individualization can be a consequence of impaired caspase activity. In *Drosophila* spermiogenesis, multiple caspase and caspase regulators act in non-apoptotic roles, to mediate spermatid individualization (Arama E et al., 2003). To determine whether *Drosophila* effector caspases become activated in Mob4^RNAi^ testes, we carried out immunostaining with the anti-cleaved caspase 3 antibody (anti-CC3) which cross-reacts with the caspase-3-like *Drosophila* effector caspase drICE. In control testes, CC3 signal was mostly restricted to the cystic bulge and accumulated in the waste bags following spermatid individualization (Figure 2B). By contrast, there were neither cystic bulges nor waste bags in Mob4^RNAi^ testes, although there was caspase activation throughout the whole length of the spermatid tail, as revealed by the CC3 positive signal (Figure 2F).

Together, these results suggest that although Mob4 is not required for caspase activation during individualization, it is necessary for synchronous migration of actin cones. In the absence of Mob4, cyst elongation occurs but disruption of the ICs prevents spermatid individualization and migration of sperm into the seminal vesicle.

### Axoneme disruption in Mob4^RNAi^ elongating cysts

To determine defects in spermiogenesis that might unlie the failure of spermatid individualization in Mob4^RNAi^ males, we examined the ultrastructure of the developing axoneme using transmission electron microscopy (TEM). During the elongation phase, the developing axoneme and major and minor mitochondrial derivatives of wild-type spermatids are encompassed by a plasma membrane, known as the the ciliary sheath (Figure 3A) (Fabrizio J et al., 1998). The major mitochondrial derivative is filled with an electron dense material of the paracrystalline body. The elongation of the two mitochondrial derivatives is required for elongation and function of the flagellum (Noguchi et al., 2011; Tates, A.D., 1971) and during this process, the minor mitochondrial derivative reduces in size and volume until individualization is completed. Several lines of evidence suggest that mitochondria are key for spermatid elongation, individualization, and protein degradation in *Drosophila melanogaster* (Noguchi T et al., 2011; Aram L et al., 2016; Cox RT et al., 2009).

**Figure 3.**
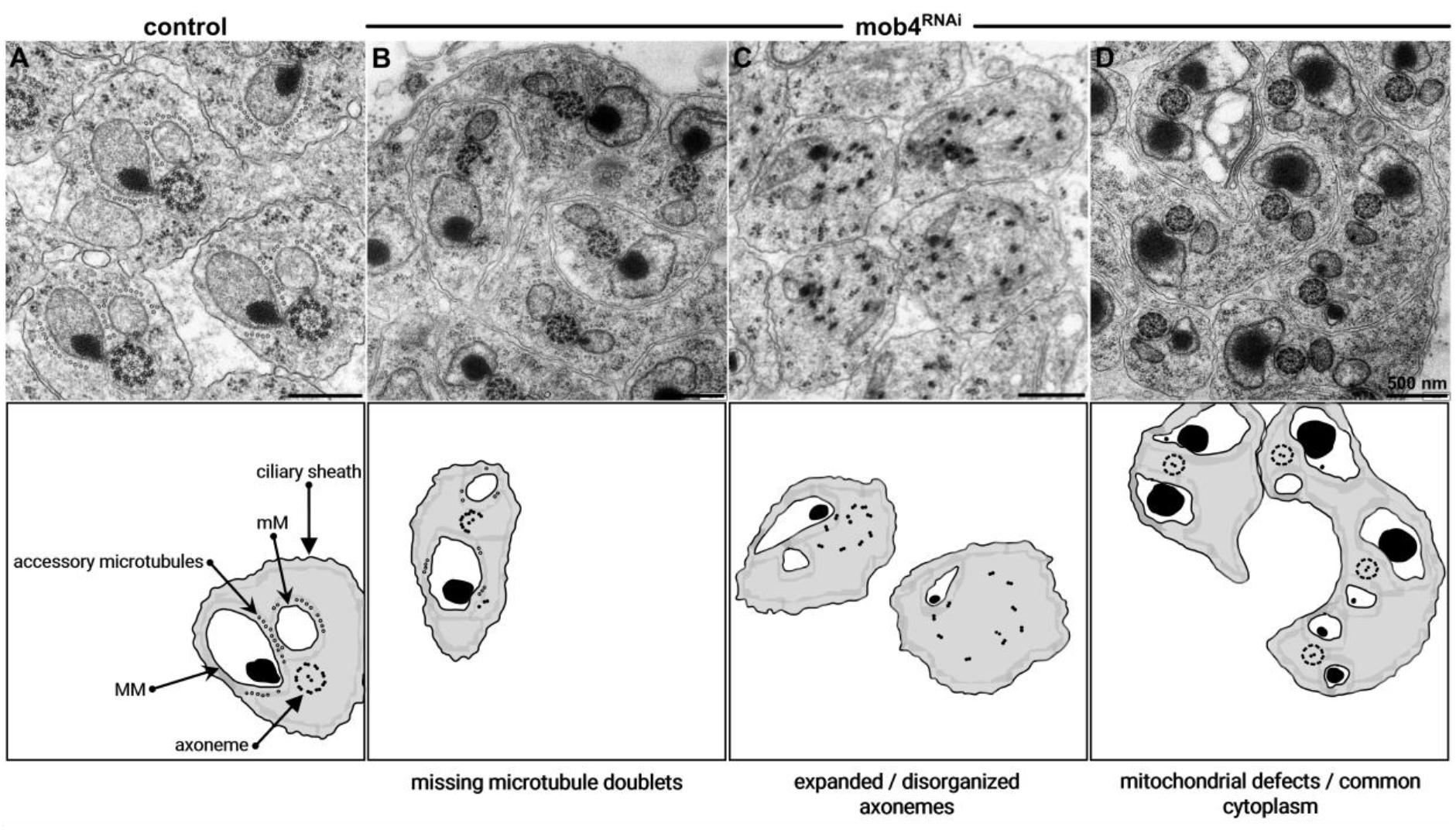
Electron microscopy reveals aberrant axonemal structure and mitochondrial defects in mob4-depleted spermatid cysts. **(A)** Transverse section of control elongating spermatids showing the major mitochondrial derivative (MM) containing paracrystalline material and the minor mitochondrial derivative (mM) adjacent to the axoneme. Accessory microtubules are in the vicinity of the mitochondrial derivatives. **(B)** mob4^RNAi^ spermatids with incomplete axonemes. **(C)** mob4^RNAi^ spermatids having “expanded” axonemes showing loss of linkage between microtubule doublets. (D) mob4^RNAi^ spermatids sharing the same ciliary sheath suggesting an incomplete second meiotic division and spermatids where both mitochondrial derivatives show a dense body.

We found that the spermatids of Mob4^RNAi^ elongating spermatid cysts displayed defects in axonemal structure before spermatid individualization was initiated. These defects included the loss of microtubule doublets thereby disrupting the 9-fold symmetry of the axoneme (Figure 3B) or preservation of stereotypical 9+2 microtubule-doublets of the axoneme but with expansion due to loss of linkage between individual doublets of the 9-fold array (Figure 3C). These findings suggest that Mob4 might not be required for axoneme formation *per se*, but for the structural maintenance of its integrity throughout the elongation stage of spermiogenesis. Moreover, spermatids with disaggregated axonemes also displayed both major and minor mitochondrial derivatives of abnormal shape and/or size. We also observed elongating Mob4^RNAi^ spermatid cysts containing two paracrystalline body-filled mitochondrial derivatives paired with one axoneme, suggesting functions of Mob4 in organizing post-meiotic mitochondria, and elongating cysts with multiple axonemes and an irregular number of mitochondrial derivatives, as occurs following failed cytokinesis (Carmena et al., 1998) (Figure 3D). Taken together, these findings suggests a role for Mob4 in the structural maintenance of axonemes and in forming or maintaining the integrity of associated cellular structures during spermatogenesis.

### GFP-Mob4 has a dynamic localization throughout spermatogenesis

Next, we examined the subcellular localization of GFP-Mob4 in the testes. Ubiquitous expression of a GFP-Mob4 transgene in the Mob4^SVC^ null mutant background rescued all phenotype, indicating that the GFP-Mob4 fusion protein is fully functional. We therefore used such transgenic flies (Mob4^SVC^;ubq-GFP-Mob4) to study the subcellular localization of GFP-Mob4 throughout spermatogenesis (Figure 4A). In meiotically dividing spermatocytes, GFP-Mob4 had a reticular localization accumulating in membranous fibers surrounding the meiotic spindle (Figure 4C). We found that during individualization, GFP-Mob4 strongly accumulated in the cystic bulge (Figure 4B) where it co-localized with cleaved-caspase 3 (data not shown). In early canoe stage spermatids, GFP-Mob4 strongly accumulated in individual punctae at the basal side of nuclei in vicinity of the basal body (Figure 4D-I). This punctate localization appeared to be very transient and was absent when the actin cone was forming in spermatids at the late canoe stage (see Supplementary Figure S1). The dynamic behaviour of Mob4 at different stages of spermatogenesis is suggestive of multiple functions in the parafusorial membranes and associated microtubules in meiosis; at the basal body or transition zone in the initiation of axoneme elongation; and in the cystic bulge during individualization *per se*.

**Figure 4.**
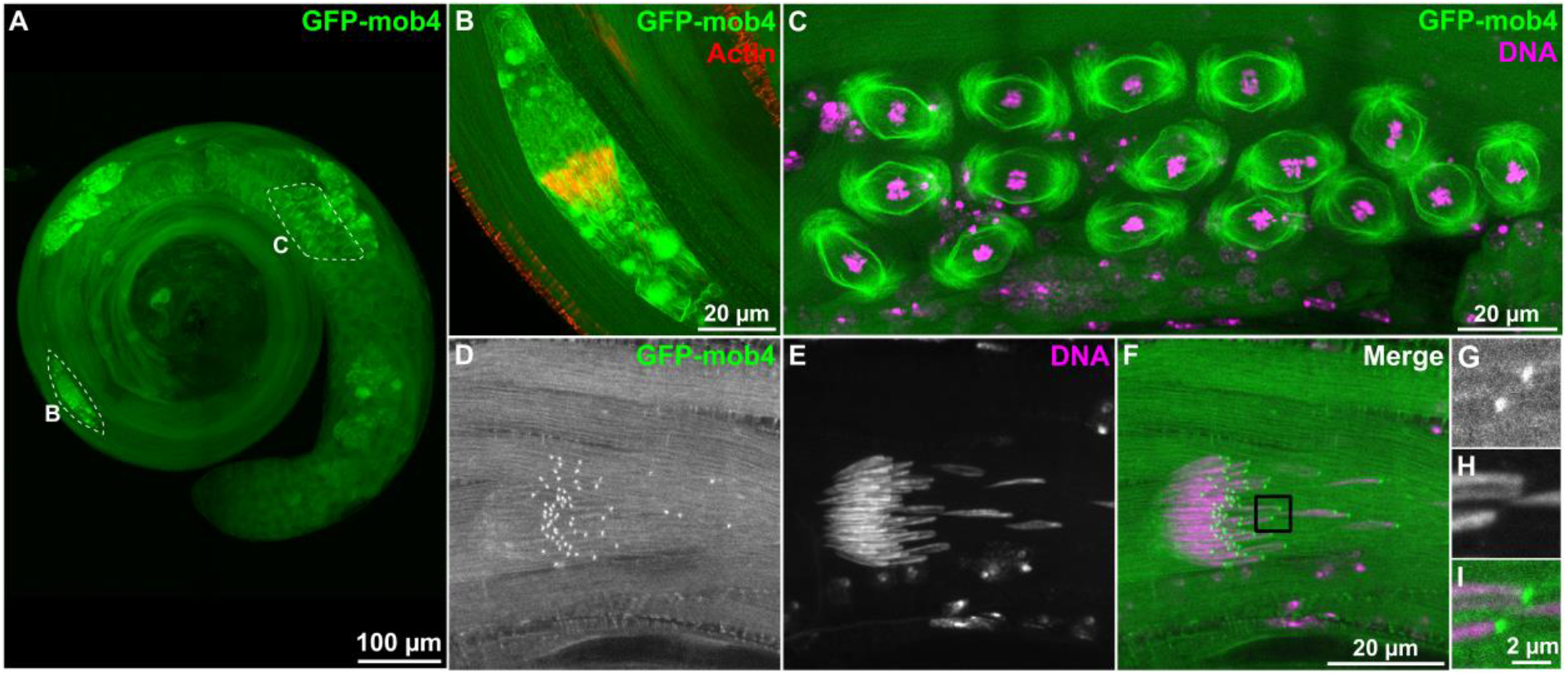
GFP-mob4 localization throughout spermatogenesis. **(A)** Testes from 2 day-old males expressing GFP-mob4 (N-terminal GFP-tagged mob4 expressed under the ubq promoter in a mob4 null mutant background) 2-day old males were stained with phalloidin-594 and DAPI to reveal actin and DNA, respectively. **(B)** GFP-mob4 accumulates in the cystic bulge surrounding actin cones in individualizing spermatid bundles. **(C)** GFP-mob4 has a reticular localization around the parafusorial membranes and associated with microtubules during the meiotic divisions. **(D-I)** GFP-mob4 localizes to the basal end of the nuclei in early canoe stage spermatids (see also Supplementary Figure S1).

### Mob4 acts through STRIPAK to support spermatogenesis and male fertility

Biochemical studies have shown that Mob4 is a core component of the macromolecular STRIPAK complex, which contains both kinase and phosphatase subunits (Goudreault M et al., 2009).

To determine whether the requirement for Mob4 in spermatogenesis reflected functions of the STRIPAK complex, we down-regulated two other core STRIPAK components, Strip and Cka, in the male germline of *Drosophila* using Strip^RNAi^ and Cka^RNAi^ RNA interference lines. As a first test of the efficacy of RNAi, we analysed the effect of ubiquitous depletion of Strip and Cka, using a da-GAL4 driver. We found that ubiquitous depletion of either Strip or Cka induced late larval/pupal lethality (Figure 5A) as previously described for alleles at these loci (Sakuma C et al., 2014; Chen HW et al., 2002). We then used nos-GAL4 and bam-GAL4 drivers to down-regulate Strip and Cka in the male germline. We found that down-regulation of Strip resulted in a male sterility phenotype comparable to that seen following down regulation of Mob4 (Figure 5B). Depletion of Cka led to strong reduction in male fertility, in which around 50% of males were completely sterile.

**Figure 5.**
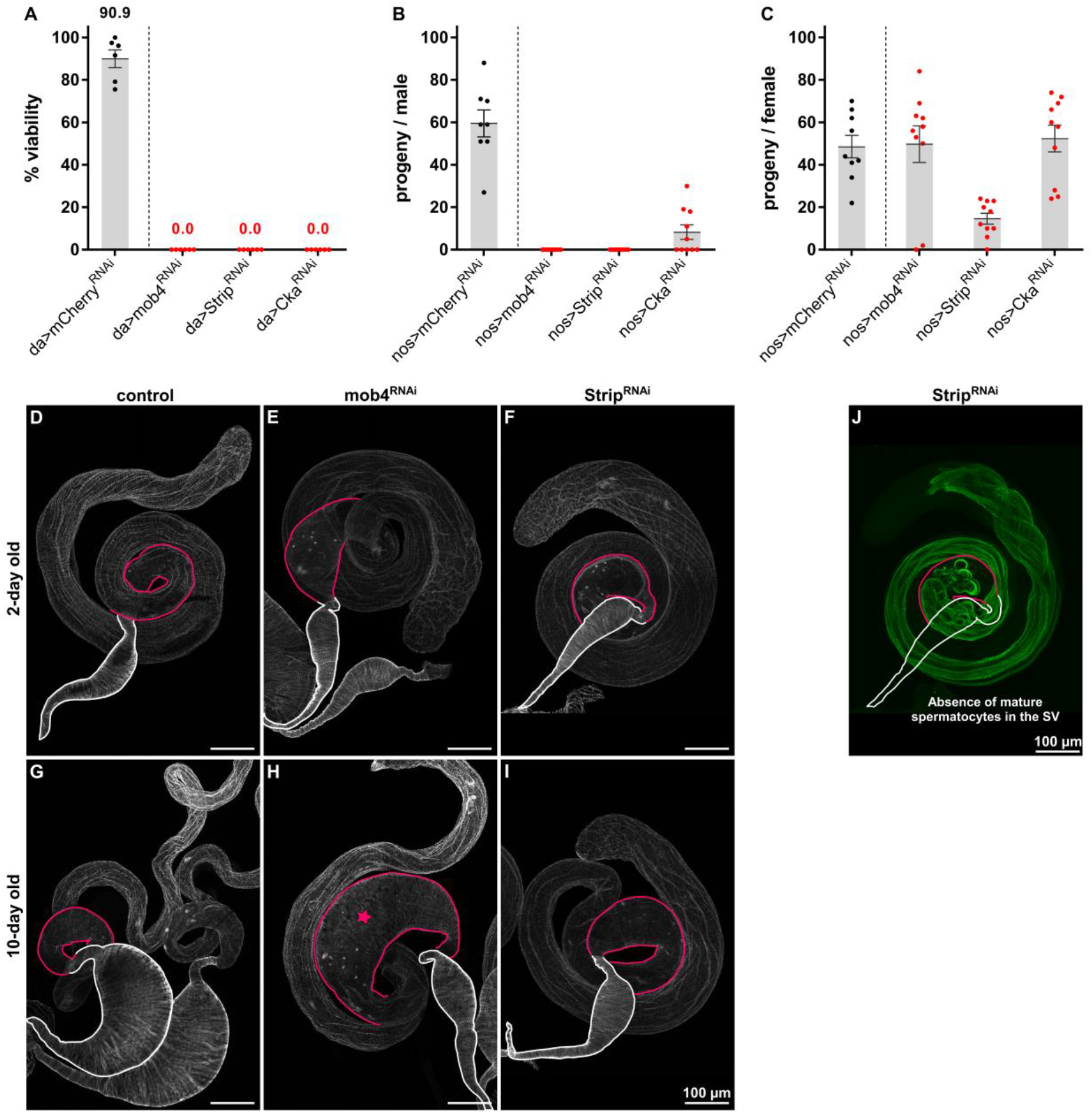
STRIPAK components Mob4, Strip and Cka are required for male fertility in *Drosophila melanogaster*. **(A)** Ubiquitous downregulation of STRIPAK core components was achieved by crossing mob4^RNAi^, Strip^RNAi^ and Cka^RNAi^ flies to the da-GAL4 driver line. Viability represents percentage of eclosed adults relative to number of pupae in 6 independent crossings. **(B)** Germinal downregulation of STRIPAK core components was achieved by crossing UAS-RNAi flies (as in A) to the nos-GAL4 driver line. Males of each genotype were individually mated with wild-type females. Data points represent numbers of progeny from individual males. **(C)** Females of each genotype were individually mated with wild-type males. Data points represent numbers of progeny from individual females. **(D-I)** Whole mount testes stained with phalloidin-594 to visualize the terminal epithelium (TE, highlighted pink) and seminal vesicles (SV, highlighted white) in 2-day old **(D-F)** and 10-day old **(G-I)** control, mob4^RNAi^ and Strip^RNAi^ un-mated male flies. At day 10, control males show large seminal vesicles due to storage of mature sperm **(G)**. Seminal vesicles of mob4^RNAi^ **(H)** and Strip^RNAi^ **(I)** 10-day old unmated males remain empty. Pink star in (H) highlights enlargement of terminal epithelium in mob4^RNAi^ males.

We next determined whether the sterility of Strip^RNAi^ males was also a consequence of an impairment in spermatid individualization as observed for Mob4^RNAi^ males. When we stained testes from 2-day old Strip^RNAi^ males with fluorescent-conjugated phalloidin, this revealed that the ICs were scattered and that no mature sperm accumulated in the seminal vesicle (Figure 5J-L), an identical phenotype to that observed in Mob4^RNAi^ testes. As the control animals aged, their seminal vesicles increased in volume, as a result of continuous sperm accumulation. However, the seminal vesicles of 10-day old Mob4^RNAi^ and Strip^RNAi^ males failed to fill with sperm and no increase in volume was observed (Figures 5G-H). Instead, the testes of Mob4-depleted males showed enlargement of the terminal epithelium (Figure 5G, pink star) suggesting continuous production and accumulation of aberrant sperm and abnormal sperm coiling. Together, this accords with a requirement for Mob4 and Strip for sperm individualization, likely as components of the STRIPAK complex.

## Discussion

Here, we describe a previously unknown function of Mob4 in the post-meiotic stages of *Drosophila melanogaster* spermiogenesis that indicates a role for Mob4 during spermatid differentiation. Male meiosis in *Drosophila* results in a cyst of 64 interconnected spermatids that undergo a complex process of elongation concomitant with formation of the sperm axoneme. The process of spermatid individualization is crucial to the formation of functional sperm. Mob4 depleted testes are able of undertaking cyst elongation after meiosis. However, spermatids fail to individualize, and seminal vesicles are devoid of mature sperm. This important function is in line with the description of the testes as the tissue of highest levels of Mob4 expression in the fly (FlyAtlas2, Krause SA et al., 2022). One of the primary defects of spermiogenesis in Mob4-depleted testes is the loss of the structural integrity of the axoneme. However, we cannot rule out that additional functions of Mob4 are also required as part of the sperm individualization process.

Spermatogenesis is an intricate developmental process, that initiates with one gonialblast, that undergoes a stereotyped series of mitotic and meiotic divisions, to generate cysts of 64 interconnected haploid spermatids. Spermiogenesis then proceeds, with cell shape changes and remodelling of all subcellular organelles from the nucleus to mitochondria, in a synchronized elongation and individualization process (Lindsley et al., 1980; Fuller M, 1993). This sequence of events requires intense cytoskeleton reorganization, not only of microtubules in axoneme elongation but also with actin having a major role in spermatid individualization (Lindsley et al., 1980; Fabrizio J et al., 1998).

Several mutants for microtubule binding proteins, have been shown to affect the axoneme structure and lead to individualization defects in the developing spermatid. Testes of *Drosophila* mutants for Bug22 (Basal body up regulated gene 22), a conserved protein that associates with basal bodies and cilia, for example, show defects in sperm individualization characterized by disrupted spermatid axonemes, that appear slightly opened or completely disassembled (Maia T et al., 2014). Flies depleted for TTLL3B glycylase, an essential member of the tubulin tyrosine ligase-like (TTLL) family, are male sterile with either missing axonemes or having axonemes composed of singlet microtubules and having sperm individualization defects (Rogowski K et al., 2009). To our knowledge, however, expanded axonemes having all nine outer doublets but with lost linkage, is a unique characteristic of Mob4 downregulation and the first time this has been observed to be associated with individualization defects.

Previous genetic studies have shown a requirement for Mob4 as a key regulator of neuronal structure and development in *Drosophila*, acting at the level of microtubule organization and axonal transport (Schulte J et al., 2010). In addition, a role for Mob4 focusing the microtubule-based mitotic spindles was also reported in cultured cells (Trammell MA et al., 2008). It was also shown that Mob4 mutants in zebrafish feature a defective microtubule network (Berger J et al., 2022). In addition, both in zebrafish and nematodes, the Mob4 protein interacts directly with the tubulin- and actin-folding TRiC/CCT chaperone complex (Berger J et al., 2022; Khabirova E et al., 2014). In the light of these findings, it is tempting to suggest that the defects in spermatogenesis that we now report in the absence of Mob4 are due to impaired microtubules functioning. Furthermore, in our own pull-downs of GFP-Mob4 from *Drosophila* embryos we identified components of the TRiC/CCT as molecular partners of GFP-Mob4 (data not shown).

Finally, Mob4 is a core component of the multimeric STRIPAK complex and indeed, we also find STRIPAK components associated with Mob4 in our pull-down experiments. Our findings of similar defects in the testes of males mutant for another STRIPAK component, Strip, accord with a role for the STRIPAK complex in regulating axonemal organization and sperm individualization.

Mitochondria are known to assist axonemal growth during spermatid elongation. Following the meiotic divisions, mitochondria associate with each developing spermatid, aggregate around the basal body and fuse into two separate derivatives that form the Nebenkern (Fuller, 1993; Anderson et al., 2009). Our TEM analysis revealed unexpected defects in the mitochondrial derivatives of the elongating cysts of Mob4RNAi males. At the onset of elongation, the two mitochondrial derivatives normally unfurl from one another and extend in parallel with the growing axoneme, providing both physical and metabolic support (Regan and Fuller, 1990). One of them accumulates a dense body of paracrystalline material and becomes the major mitochondrial derivative. The other reduces in size and volume until individualization is completed. In Mob4^RNAi^ elongating cysts, it was possible to observe cases of the accumulation of paracrystalline material in both mitochondrial derivatives. A similar phenotype has been described for the big bubble 8 (bb8) mutant, where spermatids showed accumulation of a dense body in both major and minor mitochondrial derivatives (Vedelek V, 2016). Moreover, following Mob4^RNAi^ both mitochondrial derivatives often show irregular shapes and dimensions, similar to what has been reported for the wampa (wam) mutant, where the mitochondrial derivatives are smaller than in wildtype (Bauerly E et al., 2020). Wampa is a dynein subunit and wam mutant spermatids lack the axonemal outer dynein arm, leading to loss of flagellar motility. It therefore seems likely that defects in axonemal organization could lead to the defects in mitochondria structure we observe. However, as the mitochondria provide structural support for the developing axoneme and are essential for the formation of the flagellum, defects in mitochondrial elongation could further destabilize the developing axoneme as previously suggested (Fuller, 1993; Hoyle and Raff, 1990).

In *Drosophila*, multiple caspases and caspase regulators participate in the individualization process, in non-apoptotic functions (Huh JR et al., 2004; Fabian and Brill, 2012). During IC progression, activated caspase forms a gradient from the cystic bulge to the distal cyst tail. In Mob4^RNAi^ testes, elevated levels of activated caspase can be seen throughout the whole length of the spermatid bundle, suggesting that Mob4 is not necessary for caspase activation. However, it is tempting to speculate that Mob4 might play a role in restricting activated caspases to the cystic bulge. In fact, pull-downs of GFP-Mob4 from syncytial embryos has identified the giant inhibitor of apoptosis (IAP)-like protein Bruce has as an interactor of Mob4 (data not shown). Loss of function mutants in Bruce are homozygous viable but male sterile, due to individualization defects and excessive caspase activity (Arama E et al., 2003).

Individualization defects can be a consequence of various impaired pathways (Steinhauer, 2015), and several *Drosophila* mutants have been described to show scattered IC phenotypes, such as Clathrin heavy chain (Chc4) mutants, which are defective in receptor-mediated endocytosis, neurotransmitter secretion, and spermatid individualization (Bazinet C et al., 1993). The possibility of common functions for Chc4 and Mob4 is tempting, given that Mob4 has a role in vesicular trafficking functions and shares sequence homology with the σ subunits of clathrin adaptor complexes (Baillat G et al., 2001; Bailly Y, Castets F, 2007).

In summary, our present study has identified cellular requirements for the Mob4 and Strip genes, which have essential roles in spermatogenesis and male fertility. Mob4 is necessary for proper axoneme structure and function and Mob4 mutants display a range of defects throughout multiple stages of sperm development. Understanding how Mob4 regulates the architecture of axonemal microtubules and how that affects other aspects of cellular function will be indispensable for understanding the pleiotropic manifestation of diseases, such as sterility, that arise from the dysfunction of STRIPAK genes.

## Materials and methods

### Fly stocks & husbandry

All fly stocks were maintained at 25°C and on standard cornmeal-yeast-sucrose media. Unless otherwise indicated, experiments were performed at 25ºC. w^1118^ flies were used as wild-type strain of *Drosophila melanogaster*. All RNAi lines used for the screen were obtained from BDSC (Bloomington Drosophila Stock Center, Indiana, USA) and are listed in Table S1. The specific stocks used and fly lines generated in this study are listed in Table S1.

### Tissue-specifc genetic manipulation

Early germ cell-specific gene depletion was accomplished using the bipartite GAL4/UAS system. Flies carrying the nosGAL4:VP16 (‘driver’ expressed in GSCs and early germ cells) and bamGAL4:VP16 (‘driver’ expressed in late spermatogonia and early spermatocytes) were crossed to flies carrying dsRNA under UAS control in the VALIUM20 (listed in table S1). Control flies were the progeny of the cross between the driver and UAS-mCherry^RNAi^.

### Viability assay

Ubiquitous depletion was accomplished by crossing flies carrying the da-GAL4 driver with flies carrying dsRNA against the genes of interest. Mating was allowed for 3 days and then flies were discarded. The number of pupae and the number of eclosed adults was scored and viability is expressed as the percentage of flies that eclosed from the pupae.

### Fertility assay

To test female fertility, virgin female flies (aged 1-3 days) depleted for the genes of interest (see *tissue-specific genetic manipulation*) were collected and individually mated with two wdil-type age-matched males. The crosses were kept at 25°C for 6 d, after which the adults were removed. The number of eclosed progeny in each vial was scored and statistically analysed. Male fertility was similarly tested as described; individual males depleted for the genes of interest were mated with two wdil-type virgin females. 10 individual crosses per genotype were evaluated in at least 2 independent experiments.

### Immunostaining of whole-mount testis

Testes were dissected from unmated young adults (2-day old males unless otherwise indicated) in PBS and fixed in 4% formaldehyde/PBS for 20-30 min. Samples were washed thrice in PBST for 20 min and blocked in PBST/10% SFB/10% BSA for 1 hour. The samples were incubated overnight in a humid chamber at 4°C with the following primary antibodies: mouse anti-pan polyglycylated tubulin (1:1000, Axo-49; Sigma) and anti-cleaved caspase 3 (1:500, Cell Signaling #9661). The samples were washed thrice in PBST for 20 min and incubated with appropriate secondary antibodies in a humid chamber for 2 h at room temperature, followed by three 20-min washes in PBST. DNA and actin were stained with 1 μg/mL DAPI and phalloidin (1:1000, Flash Phalloidin™ Red 594; BioLegend). Testis were washed three times with PBS and mounted in (ProLong™ Gold Antifade Mountant; Invitrogen) or Mowiol.

### Confocal microscopy

Microscopic images were acquired on a Zeiss LSM710 confocal microscope (using ZEN blue software, Zeiss) using 25× (oil) or 100× (oil) objectives. Stitching of tile scan microscope images was performed using ZEN black software (Zeiss). All processing and analysis of microscope images was performed with ImageJ (Schneider C *et al*., 2012) (ImageJ 1.53q). Figures were created using the QuickFigures2 plugin for FIJI/ImageJ (Mazo G, 2021).

## Supporting information

Mob4 Spermatogenesis Supplemental Fig.1

## Acknowledgments

Work funded by Algarve 2020 Program, grant number ALG-01-0145-FEDER-030014, and co-financed by FEDER Funds through the Operational Program for Competitiveness Factors—COMPETE 2020 and by National Funds through FCT—Foundation for Science and Technology under the Project PTDC/BIA-CEL/30014/2017. Inês B. Santos funded by FCT fellowship SFRH/BD/141734/2018. Giulliano Callaini funded by Ministero dell’Istruzione, dell’Università e della Ricerca (MIUR 2020CLZ5XW).

*Drosophila* stocks obtained from the Bloomington Drosophila Stock Center (NIH P40OD018537) were used in this study. We also thank Flybase (Gramates LS et al., 2022) and FlyAtlas (Krause SA et al., 2022) for providing database information.

## Notes

### Competing Interest Statement

The authors have declared no competing interest.

