## Supplementary material for "Mob4 is essential for spermatogenesis in *Drosophila melanogaster*": Mob4 Spermatogenesis Supplemental Fig.1

**Supplemental Figure 1**

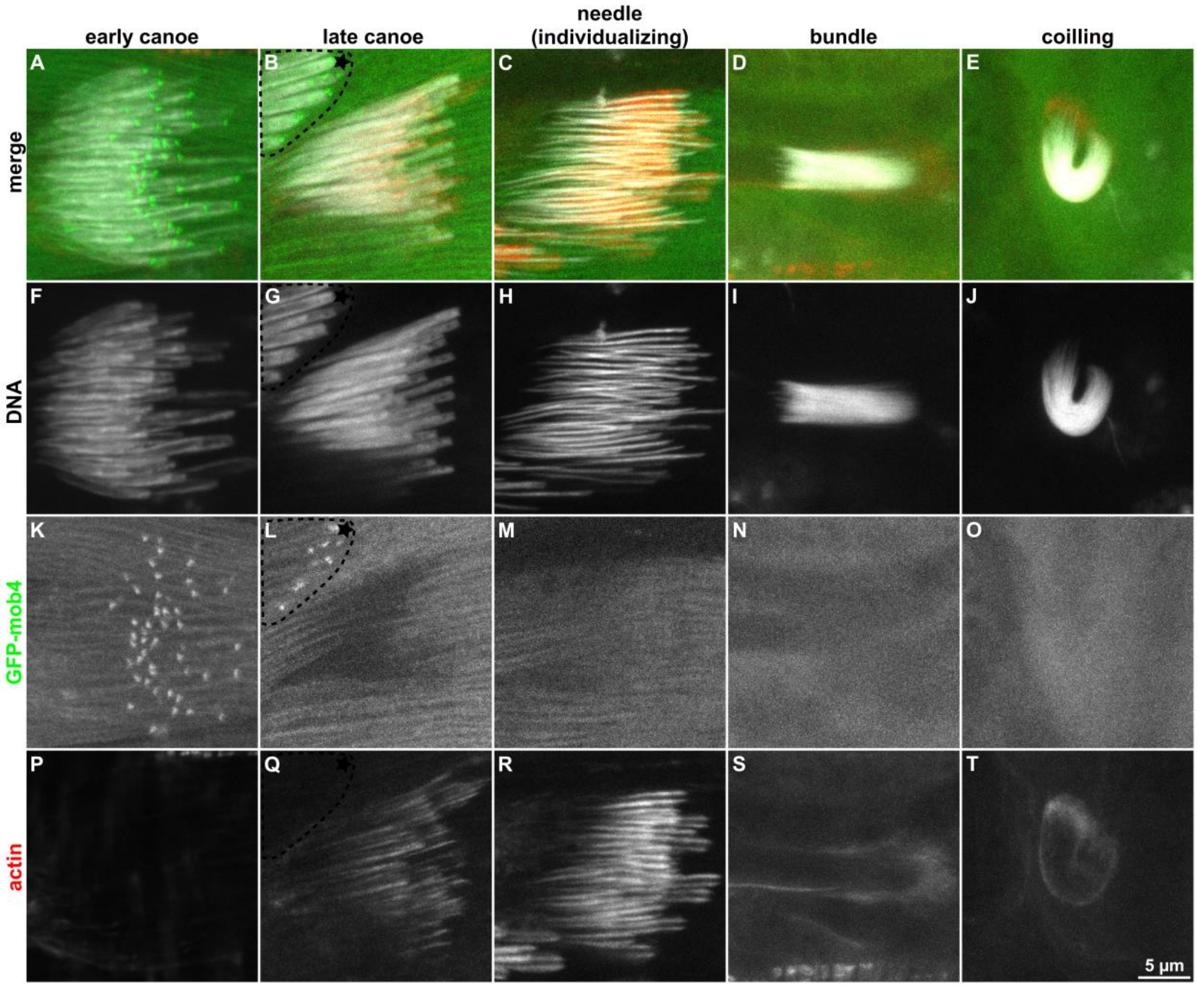


Supplementary figure S1. **GFP-mob4 transient localization at the basal body.** GFP-mob4 has a dynamic localization throughout spermiogenesis **(A-E)**. Testis from GFP-mob4 2-day old males we're visualized with DAPI **(F-J)** and phalloidin-594 **(P-T)** to reveal the localization of GFP-mob4 **(K-O)** relative to the spermatid nuclei and actin cone assembly. GFP-mob4 localizes to the basal end of the nuclei in early canoe stage spermatids **(K)** but that localization is lost after actin cone assembly in late canoe stage spermatids **(L-O)**. Nuclei marked with a star in (B,G,L,Q) belong to early canoe stage spermatids.
